# Lipidomic and Proteomic Insights from Extracellular Vesicles in Postmortem Dorsolateral Prefrontal Cortex Reveal Substance Use Disorder-Induced Brain Changes

**DOI:** 10.1101/2024.08.09.607388

**Authors:** Chioma M. Okeoma, Wasifa Naushad, Bryson C. Okeoma, Carlos Gartner, Yulica Santos-Ortega, Calvin Vary, Victor Corasolla Carregari, Martin R. Larsen, Alessio Noghero, Rodrigo Grassi-Oliveira, Consuelo Walss-Bass

## Abstract

Substance use disorder (SUD) significantly increases the risk of neurotoxicity, inflammation, oxidative stress, and impaired neuroplasticity. The activation of inflammatory pathways by substances may lead to glial activation and chronic neuroinflammation, potentially mediated by the release of extracellular particles (EPs), such as extracellular condensates (ECs) and extracellular vesicles (EVs). These particles, which reflect the physiological, pathophysiological, and metabolic states of their cells of origin, might carry molecular signatures indicative of SUD. In particular, our study investigated neuroinflammatory signatures in SUD by isolating EVs from the dorsolateral prefrontal cortex (dlPFC) Brodmann’s area 9 (BA9) in postmortem subjects. We isolated BA9-derived EVs from postmortem brain tissues of eight individuals (controls: n=4, SUD: n=4). The EVs were analyzed for physical properties (concentration, size, zeta potential, morphology) and subjected to integrative multi-omics analysis to profile the lipidomic and proteomic characteristics. We assessed the interactions and bioactivity of EVs by evaluating their uptake by glial cells. We further assessed the effects of EVs on complement mRNA expression in glial cells as well as their effects on microglial migration. No significant differences in EV concentration, size, zeta potential, or surface markers were observed between SUD and control groups. However, lipidomic analysis revealed significant enrichment of glycerophosphoinositol bisphosphate (PIP2) in SUD EVs. Proteomic analysis indicates downregulation of SERPINB12, ACYP2, CAMK1D, DSC1, and FLNB, and upregulation of C4A, C3, and ALB in SUD EVs. Gene ontology and protein-protein interactome analyses highlight functions such as cell motility, focal adhesion, and acute phase response signaling that is associated with the identified proteins. Both control and SUD EVs increased C3 and C4 mRNA expression in microglia, but only SUD EVs upregulated these genes in astrocytes. SUD EVs also significantly enhanced microglial migration in a wound healing assay.This study successfully isolated EVs from postmortem brains and used a multi-omics approach to identify EV-associated lipids and proteins in SUD. Elevated C3 and C4 in SUD EVs and the distinct effects of EVs on glial cells suggest a crucial role in acute phase response signaling and neuroinflammation.

## Introduction

The brain is safeguarded by the blood-brain barrier (BBB), a complex structure comprising a heterogeneous population of cells, including brain microvascular endothelial cells (BMVECs) and components of the neurovascular unit (astrocytes, pericytes, neurons, and the basement membrane). The interaction between BMVECs and the neurovascular unit is crucial for intercellular communication, ensuring the proper functioning of the central nervous system (CNS). In SUD, the demand for intercellular communication escalates due to the influx of inflammatory cells and particles into the brain parenchyma. Understanding the drivers of brain inflammation is essential for advancing our knowledge of CNS mechanics in SUD ^1-13^. One mechanism of cell-to-cell communication is through the release and internalization of EVs, which are membrane-bound vesicles that encapsulate proteins, lipids, nucleic acids, and other cargos. EVs are formed through processes such as plasma membrane budding and endosomal system packaging, and they are released by various cell types, including those in the CNS ^14, 15^.

In the CNS, EVs play significant roles in neuronal activation ^16^ and mediate communication between neurons and astrocytes ^17^, oligodendrocytes ^18^, and microglia ^19^. In SUD, EVs are thought to regulate responses to substances of abuse, including cocaine ^20, 21^, cannabinoids ^15, 22^ nicotine ^23^, alcohol ^24^, and opioids ^25^. However, research on EVs in humans with SUD remains limited. Most studies have utilized EVs isolated from cell lines, limiting the exploration of EV interactions with CNS cells in the brain’s intricate microenvironment ─ a situation that is further exacerbated by the difficulty in isolating EVs from brain tissues.

Our current study advances the field of SUD biology by using particle purification chromatography (PPLC) ^22, 26^ in isolating EVs from postmortem dlPFC of individuals with SUD and without SUD (controls). We assessed the EVs’ physical properties, evaluated their cargo through lipidomic and proteomic analyses, and examined their bioactivity via functional assays of cellular uptake, as well as their effects on cellular gene expression and cell migration.

## Methods

### Postmortem brain tissues

Sample demographics are described in **Table 1**. Postmortem brain tissues were obtained from the University of Texas Health Science Center at Houston (UTHealth Houston) Brain Collection in collaboration with the Harris County Institute of Forensic Science, with approval from the Institutional Review Board, and after consent from the next of kin (NOK). Medical examiner reports including cause of death and toxicology were obtained and medical records were acquired when available. For SUD subjects, substance-induced toxicity (cocaine and/or opioids) was confirmed as the cause of death by medical examiner’s report and toxicology after death. All controls died of cardiovascular disease. A detailed psychological autopsy of the donor was obtained by interviewing the NOK ^27^ from which information of age of onset of drug use, types of drugs used, smoking and drinking history, and any co-morbidities was obtained. A diagnosis of OUD or CUD, or designation as non-psychiatric control (absence of any apparent psychopathology), was determined according to DSM-5 criteria after a consensus meeting where three trained clinicians reviewed the psychological autopsy and all other available records. Upon receipt of the brain, the right hemisphere was coronally sectioned, immediately frozen, and stored at -80º C. Postmortem interval (PMI) which ranged from 13.07 and 39.35 hours was calculated from the estimated time of death until tissue preservation. Dissections of BA9, defined within the dlPFC between the superior frontal gyrus and the cingulate sulcus, were obtained using a 4 mm cortical punch.

**Table 1:** Donor demographics and sample characteristics.

| SAB | Group | Ethnicity | Gender | Age | PMI (hours) | Cause of Death | Drugs at time of death | Overdose | Diagnosis | Tissue weight (mg) |
| --- | --- | --- | --- | --- | --- | --- | --- | --- | --- | --- |
| 67951 | SUD | Black | Female | 58 | 39.35 | Toxic effect of cocaine | Cocaine, cocaethylene, benzoylecgonine, tetramisole | X | CUD | 135 |
| 67968 | SUD | Black | Female | 45 | 13.07 | Toxic effect of cocaine | Cocaine and metabolites | X | CUD | 139 |
| 67958 | SUD | White | Male | 36 | 15.05 | Toxic effects of heroin | 6-monoacetylmorphine, morphine, codeine | X | OUD | 161 |
| 67996 | SUD | White | Female | 36 | 24.47 | Toxic effects of hydrocodone and morphine | Hydrocodone and morphine | X | OUD | 126 |
| 79204 | Control | Black | Female | 45 | 26.40 | Asthma | Neg |  | Control | 119 |
| 79225 | Control | Black | Female | 69 | 20.32 | Cardiovascular disease | Neg |  | Control | 128 |
| 67976 | Control | White | Female | 69 | 24.82 | Cardiovascular disease | Neg |  | Control | 127 |
| 67992 | Control | Black | Female | 64 | 28.03 | Cardiovascular disease | Neg |  | Control | 139 |

### Isolation of postmortem BA9 EVs

The term EVs as used in this study encompasses subgroups of exosomes, microvesicles, and other membranous vesicles, which have overlapping size, density, charge, and surface markers ^28^. These subgroups cannot be accurately separated from each other on the basis of their physical properties. Thus, in this manuscript, we did not differentiate the different subgroups of EVs and we characterized them as recommended by the Minimal Information for Studies of Extracellular Vesicles (MISEV) 2018 ^29^. The isolation of EVs typically involves a combination of techniques like ultracentrifugation, immunoaffinity capture, size exclusion chromatography (SEC), or PPLC. The chosen isolation method directly impacts the purity of EVs; therefore, in this study, we used PPLC to separate EVs from other bioactive extracellular components that may co-purify with EVs. That said, we efficiently separated the EVs population from analytes that may masquerade as EVs, such as liposomes and other non-lipid non-membrane aggregates of nucleic acids and proteins, such as ECs ^14, 15^. Our research group developed a protocol for the isolation of preparative amounts of EVs from brain tissues ^22^. The schematic and workflow for isolation of EVs is shown in **Figure 1A**. Briefly, small chunks of frozen BA9 tissues, ranging from 119 mg -161 mg (**Table 1)**, were finely chopped and digested with collagenase III^22^. Samples were clarified by successive centrifugations at 500xg, 2,500xg, and then 12,00xg to remove all the cells and cell debris prior to isolation of EVs. The clarified samples were loaded on a 20 × 0.5cm Sephadex G-50 size exclusion column and purified using PPLC system as previously described^22, 26^. Fifty fractions of 200 μL were collected, and the 3D UV-Vis (230 – 650nm) fractionation profiles were recorded and used to identify fractions 8 to 21 as EV-containing fractions ^26^. A no-tissue collagenase III control was used as background. After background subtraction and PPLC analysis, EV-containing fractions were pooled, aliquoted, and stored in small aliquots at -80°C until needed for analysis.

**Figure 1:**
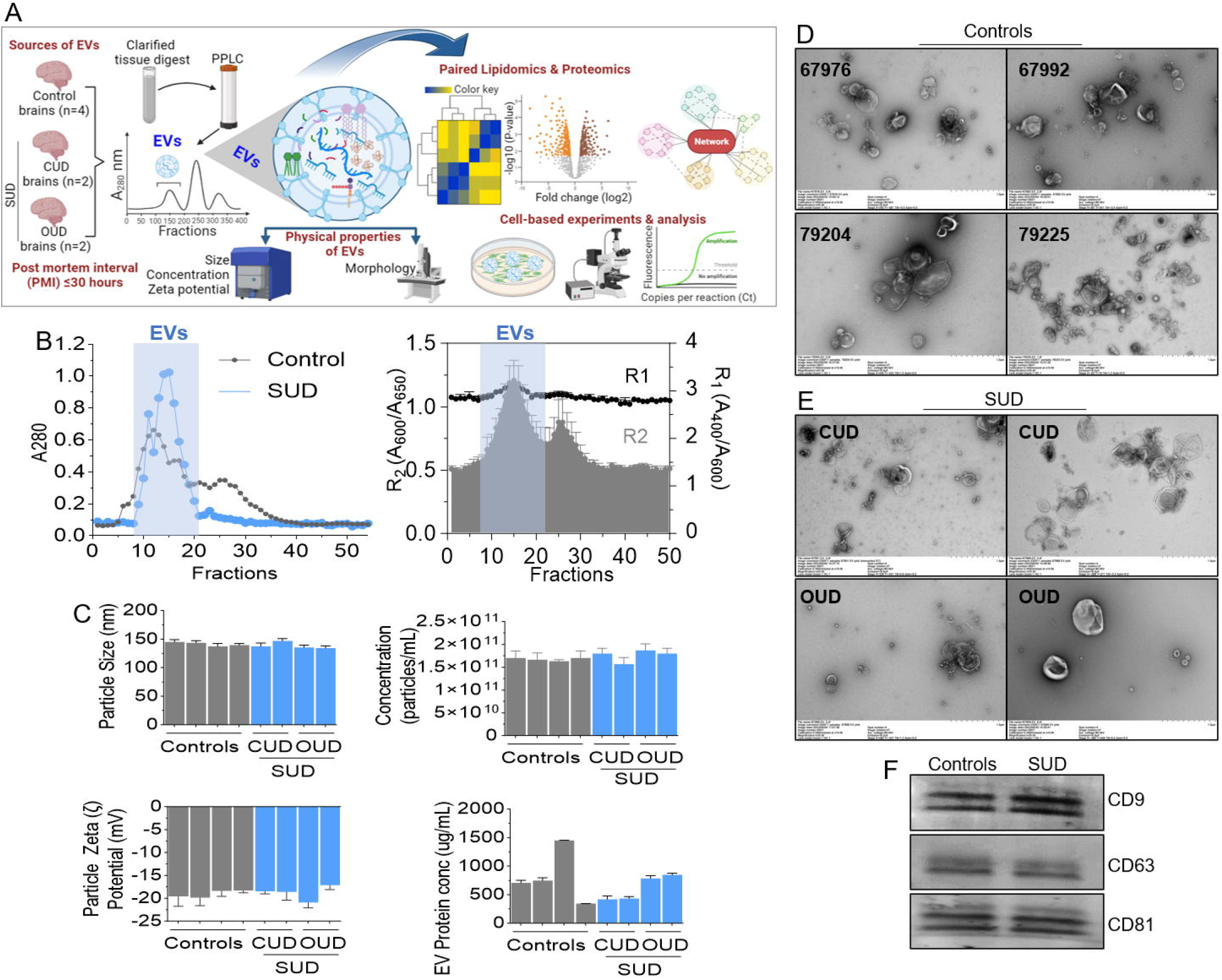
Physicochemical characterization of Postmortem BA9 EVs from control and SUD individuals. A) Infographic of experimental reagents, tools, and protocol. B) Absorbance at 280 nm of PPLC-isolated collagenase-digested BA9 eluates (left) and PPLC R2 index detected in fractions 8-21, confirming the presence of EVs in these fractions from control and SUD groups (n=4/group). C) Size (left, top), concentration (right, top), Zeta (ζ)-potential (left, bottom) of BA9 EVs measured by NTA (ZetaView), and total protein (right, bottom) measured by the Bradford assay. N=4 per group for control and n=4 for SUD (CUD, n=2, OUD, n=2). D) Representative TEM images of BA9 EVs from control and SUD individuals. E) Western blot of showing tetraspanin – CD9, CD63, and CD81 EV markers present in BA9 EVs. Error bar represent standard error of the mean.

### Assessment of EV concentration, size distribution, and surface charge

To further these analyses, EVs were diluted in 0.1X PBS (1/1000). EV size, concentration, and zeta potential (ζ-potential) measurements were acquired using nanoparticle tracking analysis (ZetaView) as described previously^22, 30^. Briefly, EV concentration (relative abundance), size distribution, and zeta potential were acquired using nanoparticle tracking analysis (NTA) (ZetaView PMX110, v8.04.02, Particle Metrix, Mebane, NC, USA) as previously described ^14, 15, 22, 26, 30^. The shutter was kept at 70, and sensitivity was adjusted to 2-4 points below the noise level to also capture small particles.

### Transmission Electron Microscope (TEM)

This is an imaging method that facilitates assessment of the purity, size, and morphology of EVs. Equal volumes of BA9 EVs from each group were pooled (n=4, n=2, n=2 for control, CUD, and OUD respectively). 10μL were spotted onto TEM grids and imaged with a Hitachi HT7700 microscope equipped with an AMT XR16M camera. Images were analyzed as described previously by us^7, 22, 26, 31^.

### Western blot for markers of BA9 EVs

Detection of BA9 EVs markers by western blot was conducted using previously published protocols^32-34^. Briefly, equal (50 μg) of BA9 EVs protein from both control and SUD groups were loaded into 4-20% (gradient) Mini-PROTEAN TGX precast gels and resolved by SDS-PAGE at a constant 100V. The separated proteins were blotted onto PVDF membranes and membranes were blocked with 5% milk dissolved in 1x TBST (50 mM Tris, 150 mM NaCl, and 0.1% Tween, pH 7.6) buffer for 1 hour at room temperature on slow shaker. The membranes were incubated at 4 °C overnight with primary antibodies against CD9, CD63, CD81. After 2 10-minute washes with 1x TBST, membranes were incubated with appropriate IRDye secondary antibodies at room temperature on slow shaker for 1.5 hours. The membranes were washed in 1X TBST for 5 minutes prior to band detection using the Li-COR Odyssey Infrared Imaging System.

### Lipid extraction

Samples were concentrated to obtain approximately10 μL for lipids extraction. Lipids were isolated using a modified Bligh and Dyer protocol ^35^ using dichloromethane and methanol. Briefly, 10 μL of each sample were transferred to a glass screw-cap tube with 90 μL of water (Burdick and Jackson-Honeywell, USA) LC-MS grade and kept on ice for 10 min. (MeOH) (Alfa Aesar-Themo Fisher, USA). Then, 2 mL of Methanol MeOH and 0.9 mL of dichloromethane (DCM) (Sigma Aldrich, USA) were added. After vortexing the DCM-MeOH-sample monophase was formed. Samples were left for 30 minutes at room temperature and aqueous and organic phases were separated by subsequent addition of 1 mL of water and 0.9 mL of DCM. All samples were centrifuged at 1200 rpm for 10 minutes and the lower DCM organic phase was removed to a new glass tube and dried under nitrogen and a partial vacuum using a Visiprep manifold (Supelco, Bellefonte, PA). Lipid extracts were dissolved in lipid load solution: MeOH/DCM (50:50, v/v) containing 10 mM ammonium acetate (NH^4^Ac) LC/MS grade (Fisher Chemical, USA) for MS analysis.

### Lipidomics

Samples were processed following our protocol for liquid, as previously described ^36^ and then infused into a QTRAP 4000 mass spectrometer. Samples were analyzed using multiple precursor ion scanning (MPIS) ^37^ in Positive and Negative polarities using the following parameters: Fragments File = Fragments_pos, Mass Tolerance (Da) = 0.3, Fragments_neg, Mass Tolerance (Da) = 0.5 spectrum peak. Direct Infusion. Types in Positive: <u>Glycerophospholipids</u> (PC, PE) total Double Bonds: From 0 To 6, <u>Sphingolipids</u> (SM, Cer, GM3, GM2, GM1, GD3, GD2, GD1) total Double Bonds: From 0 To 4. <u>Glycerolipids</u> (DAG, TAG) total Double Bonds: From 0 To 12.Types in Negative: <u>Glycerophospholipids</u> (PA, PC, PE, PG, PI, PS) total Double Bonds: From 0 To 12. Lipidomics data were processed using LipidView software (Version: 1.3 beta) and Marker View (version 1.4); Normalized using <u>M</u>ost <u>L</u>ikely <u>R</u>atio (MLR) Method, Welch t-Test and Principal component analysis (PCA) using the Pareto-scaled method.

### Proteomics

Protein data-independent quantitation (e.g., SWATH) and data-dependent ion spectra libraries were generated as previously describe d ^38-40^. Protein identification was determined using the Paragon algorithm ^41^ in the ProteinPilot (SCIEX) software package, with a <5% False Discovery Rate (FDR). For quantitative analysis, data independent mass spectrometric acquisition using Variable Window SWATH Acquisition ^42^ was conducted in triplicate and used to determine optimal Q1 isolation windows and increase specificity of detected ions. Peaks were extracted with 95% peptide confidence and 1% FDR. Quantification of SWATH and MRM data acquisition was accomplished using in MarkerView ^43^ (Sciex) and normalized using MLR normalization ^44^ for SWATH and total area sum normalization for MRM, as described previously ^45, 46^. P-values for fold change calculated using Welch’s t-test in MarkerView (Sciex).

### Lipidomic and Proteomic Data Analysis

Lipidomics and proteomics data were analyzed using the multiple precursor ion scanning (MPIS) workflow ^47^, LipidView software (Version: 1.3 beta) and Marker View (version 1.4) packages, as previously described ^48, 49^. Lipidomics data were analyzed following normalization based on common endogenous lipid classes and total lipid using the “Most Likely Ratio” method ^44, 50^. Principle component analysis of data was conducted as previously described ^43, 51^. Data set group comparisons were analyzed using unequal variance (Welch’s) t-test unless otherwise indicated ^43^. A p-value of less than 0.05 was considered significant.

### Labeling of BA9 EVs and cellular uptake assay

Pooled BA9 EVs (50 μg) were concentrated using amicon ultra-0.5 centrifugal filter units (Millipore Sigma) by centrifuging at 14,000g for 5 minutes to remove excess PBS. Resulting volumes were filled to 100 μl to label EVs with green, fluorescent dye (SYTO™ RNASelect™ stain for RNA - Invitrogen™) according to the manufacturer’s instructions. After labeling, unincorporated dye was removed using exosome spin columns (MW3000 - Invitrogen™) according to the manufacturer’s instructions. Labeled PBS was used as negative control. 100,000 cells/100 μL of primary human astrocytes and HMC3 microglia cells seeded in each well (3 wells per 3 independent experiments) of a 96-well plate were treated with labeled BA9 EVs at 50 μg /mL and incubated for 6 hours and 48 hours. Images of cells with internalized fluorescently labeled EVs were obtained after 6 and 24 hours using BioTek Lionheart FX automated microscope at 10x. Fluorescence intensity was computed using Gen5 software and plotted using Prism GraphPad version 10.

### Isolation of RNA from glia cells treated with BA9 EVs and real-time quantitative PCR

Equivalent numbers (100,000) of human primary astrocytes and HMC3 microglia cells were treated with 50 μg/mL of BA9 EVs for 24 hours in triplicates. RNA was isolated using Zymogen Quick RNA isolation kit, per manufacturer’s protocol. RNA was eluted and the eluate was measured using a NanoDrop 1000. 1 μg of total RNA was used for cDNA synthesis. 500 μg of cDNA was used for analysis of gene expression by real-time quantitative PCR (RT-qPCR) that was performed using a 7500 FAST machine and Power Track SYBR Green master mix as previously described. Primers used for the RT-qPCR are listed on **Table 2**.

**Table 2:**
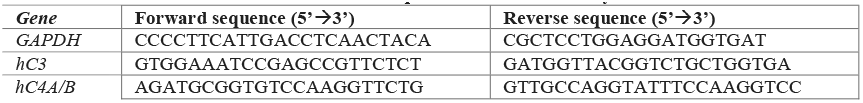
Primer sequences used in this study.

### Flow Cytometry

Equivalent concentration (200 μg/mL) of control and SUD BA9 EVs were resuspended in 0.1X PBS. Unlabeled Dynabeads™ in PBS were used as negative control. Clarified culture supernatants from confluent monolayers of human primary astrocytes and HMC3 microglia cultured with EV-depleted FBS were used as positive controls. 20 μg of biotinylated antibodies for specific cell surface markers, including GFAP, GLAST-1 for astrocytes and IBA-1 for microglia were coupled to streptavidin Dynabeads™. The antibody-bead complex was added to 200 μg/mL BA9 EVs and the mixture incubated overnight at 4 °C with gentle mixing. The next day, captured BA9 EVs were magnetically separated, washed three times, and then resuspended in 500 μL MACSQuant running buffer (Miltenyi Biotec). Flow cytometry was performed using BD FACSCelesta™ flow cytometer. Flow cytometry data analysis was performed using FlowJo™ v10.10.

### Cell migration analysis using wound healing (scratch) assay

To perform the cell migration assay, we used our previoulsy published wound healing assay protocol^52^. Briefly, 50,000 HC69 human microglia cells were seeded in silicone cell culture inserts (2 well in μ-Dish 35) mm with a defined cell-free gap within the monolayer (Ibidi), designed for precise cell migration (wound healing) assay. The cells were cultured for 5 hours and at this time, the cells were adherent to the dish. 50 μg of control or SUD BA9 EVs were added to the cells. 18 hours later, cells were washed and fed with serum free media (starvation) for 2 hours. Migration of cells into the wound area (gap) was monitored by imaging at different time points using BioTek Lionheart FX automated microscope. Images were analyzed to ascertain the wound closure rate (a typical experimental readout), by using BioTek Gen5 Software.

### Statistical analysis

Ingenuity Pathways Analysis (IPA) web-based bioinformatics inbuilt application was used for analysis and integration of protein to function data. The significance cutoff was set to fold change (FC) >2.0 or <-2.0 and a p-value <0.05. Ordinary One-way ANOVA or multiple comparison with Šídák’s multiple comparisons test or two-way ANOVA with Dunnett’s corrections were used to assess statistical differences. When stated, unpaired T-test with Welch’s correction was also used. Details of specific statistics are presented in each figure legend where the p-values are listed. Label-free quantitative analysis was obtained using the relative abundance intensity normalized by the median, using all peptides identified for normalization. The expression analysis was performed considering technical replicates available for each experimental condition following the hypothesis that each group is an independent variable. It was considered only proteins that were present in 2 out of three technical replicates and a statistical cutoff of ANOVA >0.05 was adopted. Protein Interaction and Pathways enrichment analyses were performed in Cytoscape and in silico analyses in an R environment ^53^. For PCA and volcano plot analysis, the MetaboAnalyst was used ^54^.

## Results

### SUD does not alter BA9 EVs physicochemical properties

Schematics for BA9 EVs isolation and subsequent analyses are shown in **Figure 1A**. The EV profiles from controls and SUD were similar in regard to size, concentration, ζ-potential (mV), and total protein concentration (**Figure 1B, C**). TEM analysis showed that control and SUD EVs (∼100-300 nm) have similar morphology and are heterogenous in size and electron density (**Figure 1D, IE)**. Further analysis showed the presence of tetraspanin markers (CD9, CD63, CD81) in both controls and SUD (**Figure 1F**). Given the small sample size, it is unknown whether SUD will alter EV morphology or if other EVs markers are present and to what extent such markers are enriched. Nevertheless, the presence and purity of the BA9 EVs preparation via the rigorous PPLC, devoid of most contaminants was demonstrated.

### Effect of SUD on EV lipid profile

We used an untargeted LC-MS/MS platform that utilizes positive/negative polarity switching to perform unbiased data dependent acquisitions (DDA) through higher energy collisional dissociation (HCD) fragmentation to profile lipid ions associated with EVs. Using the Lipid Ontology (LION) enrichment analysis web application (LION/web), a total of 98 positive (**Supplemental Table 1**) and 385 negative (**Supplemental Table 2**) polarity lipid species were identified as displayed in the heatmaps (**Supplemental Figure 1A, B**), highlighting the enrichment, and clustering of various lipid species in control and SUD. The identified EV -associated lipids were grouped into 3 main categories including sphingolipids, neutral lipids, and phospholipids that were equally distributed in control and SUD (**Figure 2A**). Additionally, the 3 lipid categories contain 16 classes of lipid (Cer, CerP, DAG, DES, GD2, GD3, GM1, GM2, GM3, GT3, Hex2Cer, Hex3Cer, HerCer, IPC, SM, TAG) in the positive polar lipids (**Figure 2B, top**) and 11 classes of lipids (CDPDAG, CL, PA, PC, PE, PG, PI, PIP, PIP2, PIP3, PS) in the negative polar lipids (**Figure 2B, bottom**).

**Figure 2:**
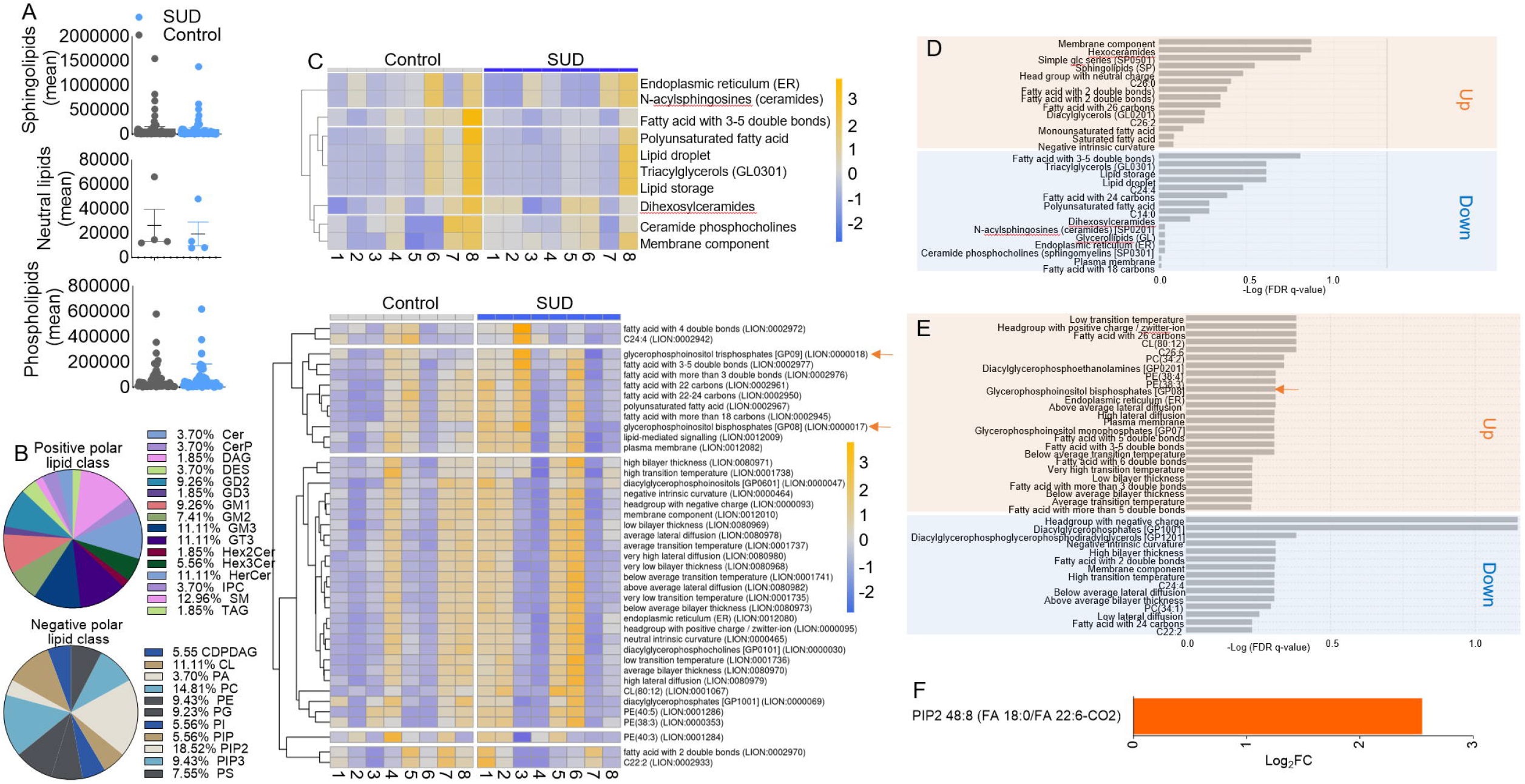
Profile of extracellular lipid classes associated with BA9 EVs from control and SUD postmortem BA9 brain tissues. **A**) Main lipid categories associated with BA9 EVs. **B**) Main lipid classes associated with BA9 EVs within the 3 lipid categories. **C)** LION-PCA heatmap for positive (Top) and negative (Bottom) polar lipids associated with BA9 EVs. **D, E**) LION enrichment analysis of BA9 EVs for **D**) positive and **E**) negative polar lipids associated with BA9 EVs. **F**) Glycerophosphoinositol bisphosphate PIP2 levels in BA9 EVs.

### LION-associations reveal differential enrichment of lipid species between control and SUD

Next, we used LION-PCA heatmap analysis to further dissect the lipidomic data). We identified significant LION-terms (**Supplemental Figure 1C, left**) or the cumulative variance explained (**Supplemental Figure 1C, right**), per set number of principal components (5) for positive ((**Supplemental Figure 1C, Top**,) and negative (**Supplemental Figure 1C, Bottom**) polar lipids, and identified major differences in EV-associated lipids between SUD and controls for positive (**Figure 2C, top**) and negative (**Figure 2C, bottom**) polar lipids. Inspection of the heatmap revealed remarkable increase in the LION-signatures of glycerophosphoinositol trisphosphates [GP09] (LION:0000018) and glycerophosphoinositol bisphosphates [GP08] (LION:0000017) in SUD BA9 EVs (**Figure 2C, bottom, orange arrows**). LION enrichment analysis identified positive and negative polar lipids that were either up or down regulated in SUD (**Figures 2D, 2E**), where Glycerophosphoinositol bisphosphate PIP2 [GP08] (LION:0000017) was strongly upregulated (**Figures 2E, orange arrow, Figure 2F**).

### Effect of SUD on EV protein profile

We used shotgun mass-spectrometry based proteomics (LC-MSMS) to profile the proteome of a portion of the same BA9 EVs ^55^ that were analyzed for their lipid content in **Figure 2**. A total of 472 proteins were identified (**Supplemental Table 3**) of which 78 were found statically dysregulated after the removal of contaminants, such as keratins (p value=0.05, FC cutoff=20% variance). A hierarchical cluster analysis applying Pearson’s correlation based on the protein expression levels demonstrates a discernible disparity between the SUD and Controls, confirming the differences in the proteome cargo of the vesicles (**Figure 3A, B, Supplemental Figure 2A**). For enrichment analysis only proteins statistically differentially regulated (pooled p-value of ≤ 0.05 and adjusted p-value (FDR) < 0.05) was considered, where 33 proteins were significantly increased, and 45 were decreased (**Figure 3C**). Gene Ontology analysis revealed a pronounced abundance of proteins originating from exosomes, vesicles, and the extracellular space, affirming a successful enrichment of vesicles derived from brain tissue (**Figure 3D**). A substantial proportion of these proteins were implicated in inflammatory and innate immune responses, (**Supplemental Figure 2B**). Biological and WikiPathways analysis indicates that the proteins may be involved cellular movement (**Figure 3E top**) and focal adhesion (**Figure 3E, bottom**). Using ingenuity pathway analysis (IPA) we identified the top 5 biological processes (**Table 3**), with infectious diseases, organismal Injury and abnormalities, metabolic disease, neurological disease, psychological disorders, as the top 5 diseases (**Table 3**). Signaling pathway analysis revealed that proteins associated with SUD EVs have the potential to affect two major cellular processes – cell viability and cell motility (**Figure 3F, supplemental Figure 2C**). The upstream regulators of cellular functions were determined by IPA for cell death (**Figure 3G, left**) and cell migration (**Figure 3G, right**). Protein to function relationships analysis of the network molecules identified human ALB, C3, and C4A/C4B proteins, all upregulated in SUD, as major hub proteins for proinflammatory cytokines (**Figure 3H, supplemental Figure 2D**). Additionally, C3 and C4A/C4B proteins are hub proteins for multiple neurodegenerative disease (**Figure 3I**).

**Table 3:** Top 5 Biological Processes associated with SUD BA9 EVs.

| Top 5 Diseases and Disorders |  |
| --- | --- |
| Name | p-value range |
| Infectious Diseases | 7.40E-03 - 1.05E-13 |
| Organismal Injury and Abnormalities | 9.24E-03 - 1.05E-13 |
| Metabolic Disease | 9.24E-03 - 1.52E-13 |
| Neurological Disease | 7.59E-03 - 1.11E-12 |
| Psychological Disorders | 7.22E-03 - 2.30E-12 |
| Top 5 Canonical Pathways |  |
| Name | p-value |
| Response to elevated platelet cytosolic Ca2+ | 4.34E-14 |
| Neutrophil degranulation | 9.86E-14 |
| Acute Phase Response Signaling | 1.31E-12 |
| LXR/RXR Activation | 1.04E-07 |
| DHCR24 Signaling Pathway | 1.97E-07 |
| Top 5 Causal Network |  |
| Name | p-value range |
| MYH9 | 3.81E-17 |
| BHLHB | 8.93E-15 |
| Mn(III)-tetrakis-(4-benzoic acid) porphyrin | 1.79E-14 |
| dimethyl fumarate | 1.96E-14 |
| tert-butyl-hydroquinone | 2.72E-14 |
| Top 5 Molecular and Cellular Functions |  |
| Name | p-value range |
| Cell Death and Survival | 7.40E-03 - 1.54E-07 |
| Cellular Movement | 9.06E-03 - 1.33E-06 |
| Cellular Assembly and Organization | 9.23E-03 - 2.82E-06 |
| Amino Acid Metabolism | 5.55E-03 - 3.36E-06 |
| Energy Production | 6.49E-03 - 3.36E-06 |
| Top 5 Physiological System Development and Function |  |
| Name | p-value range |
| Cardiovascular System Development and Function | 9.24E-03 - 5.58E-07 |
| Organismal Development | 9.24E-03 - 5.58E-07 |
| Tissue Development | 9.24E-03 - 2.82E-06 |
| Hematological Sys Development & Fun. | 9.24E-03 - 1.27E-05 |
| Immune Cell Trafficking | 9.00E-03 - 1.64E-05 |

**Figure 3:**
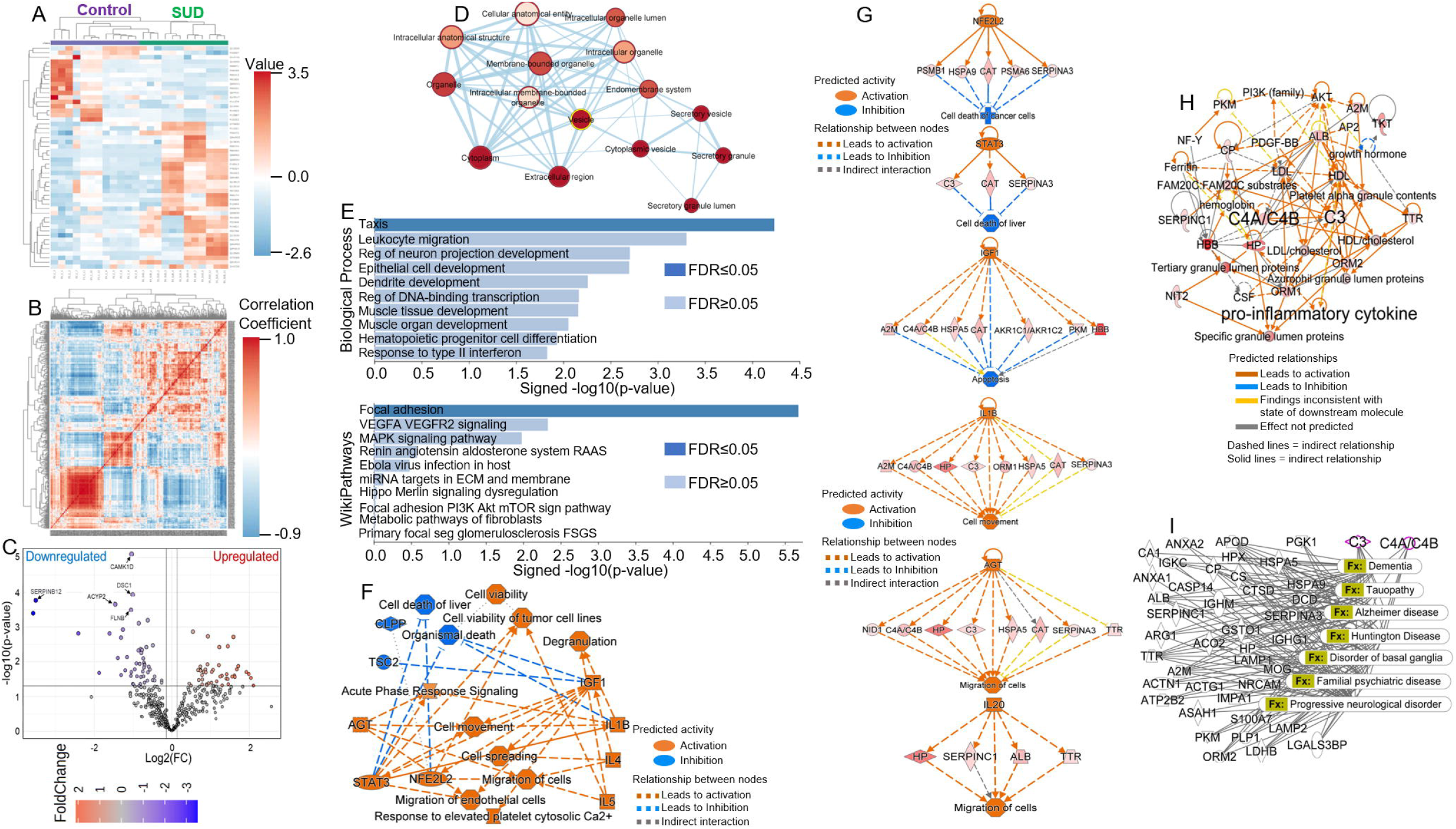
Analyses of the proteomes of BA9 EVs from control and SUD BA9 brain tissues. **A, B**) Hierarchical cluster analysis assessing differences in BA9 EVs from control and SUD BA9 tissues using Pearson’s correlation reveals differences visualized as **A**) heatmap and **B**) correlation. **C**) Volcano plot analysis of BA9 proteome. **D**) Gene Ontology analysis showing abundance of proteins originating from exosomes, vesicles, and the extracellular space, affirming a successful enrichment of vesicles derived from brain tissue. **E)** Gene Ontology analysis of BA9 proteome using WEB-based GEne SeT AnaLysis Toolkit (WebGestalt 2024). **F**) Ingenuity pathway analysis (IPA) predicted ontologically-related functions and molecules predicted to be increased (orange) or decreased (blue) driven by SUD induced changes in proteome of BA9 EVs. Functions are biological processes like cell viability, cell migration, degranulation, that are not themselves diseases, while molecules are IL1B, AGT, STAT3 and others that may regulate the functions. A solid line represents a direct interaction between the two gene products and a dotted line means there is an indirect interaction. **G**) IPA-identified upstream regulators of cellular functions for cell death (left) and cell migration (right) mediated by BA9 EVs. A solid line represents a direct interaction between the two gene products and a dotted line means there is an indirect interaction. Orange is activation or increased, blue is inhibition or decrease. **H**) IPA-identified most highly rated network and canonical pathway analysis. **I**) IPA-identified groups of ontologically-related diseases predicted to be linked to the proteome of BA9 EVs.

### SUD regulates the release of EVs by microglia cells

The majority of EVs were positive for the microglia marker IBA 1 (**Figure 4A**), suggesting that EVs were predominantly of microglia rather than astrocyte origin. Additional analysis comparing control versus SUD revealed that microglia within the dlPFC may shed significantly more EVs in response to SUD compared with control (**Figures 4B, 4C**).

**Figure 4:**
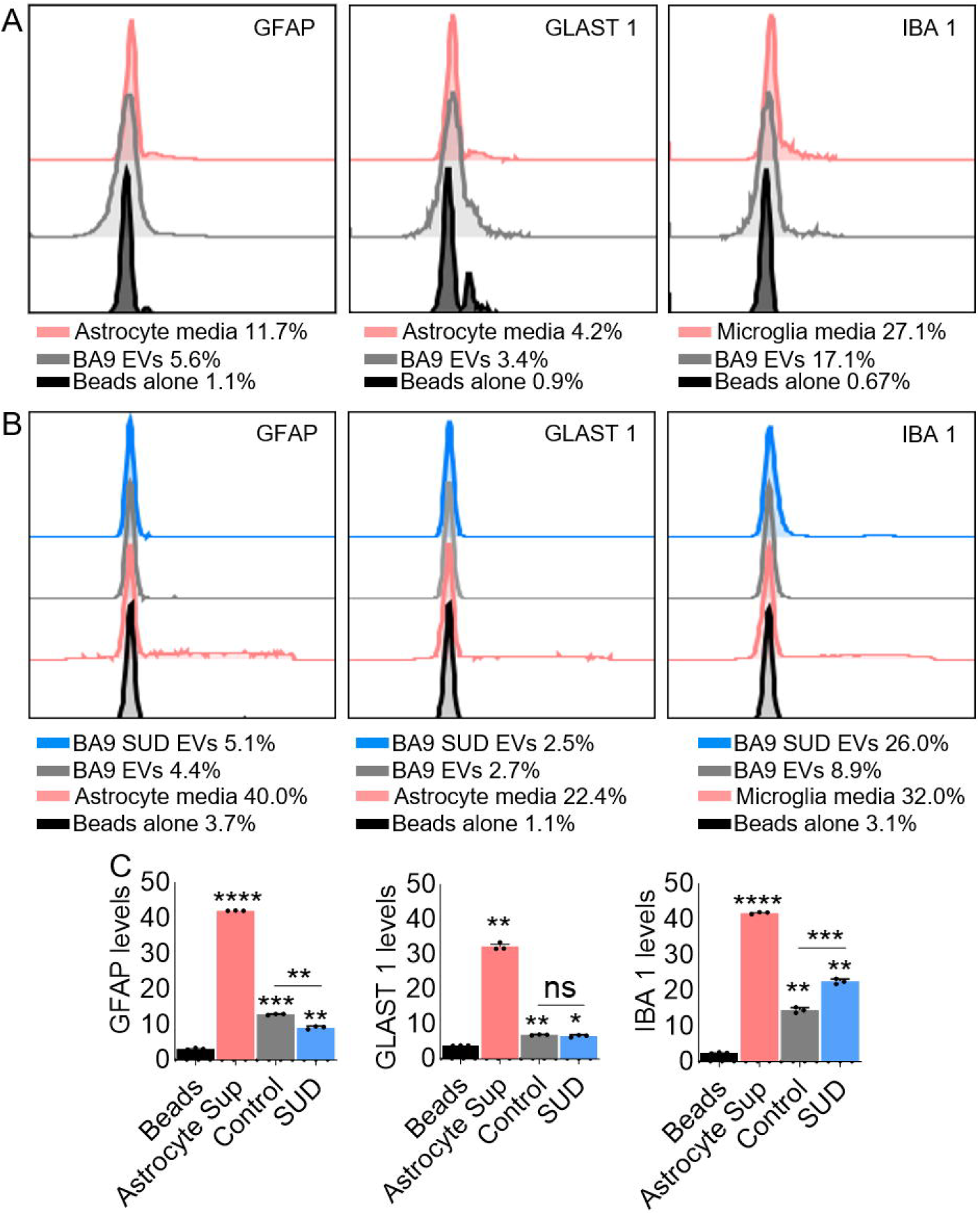
Flow cytometry analysis of BA9 EVs isolated from postmortem brains of control and SUD individuals. **A**) Pooled control and SUD BA9 EVs demonstrating presence of glia cell derived EVs irrespective of substance use. **B**, **C**) Control and SUD BA9 EVs showing differences in the release of EVs by control and SUD BA9 postmortem brain tissues as visualized by **B**) histograms and **C**) quantitative values presented in bar charts. EVs isolated from astrocyte or microglia media were used as positive control, while beads alone served as negative control. Statistical significance was determined by ordinary one-way ANOVA (Šídák’s multiple comparisons test) and unpaired t test with Welch’s correction. **** p < 0.0001, *** p 0.0005, ** p 0.0088 - 0.0051, * p 0.0167, ns = non-significant.

### Glia cells internalize EVs

Internalization of labeled control and SUD EVs by human primary astrocytes and HMC3 microglia cells revealed green signal (**Figure 5A, 5B**) that increased over time (**Figure 5C, 5D**). At 6 hours post treatment, internalization of control EVs was significantly higher compared to SUD in astrocytes (**Figure 5C)** but not microglia (**Figure 5D**). The reason for the delayed uptake of EVs by astrocytes is unknown, but by 24 hours post treatment, there was no significant differences in uptake of EVs by the two cell types.

**Figure 5:**
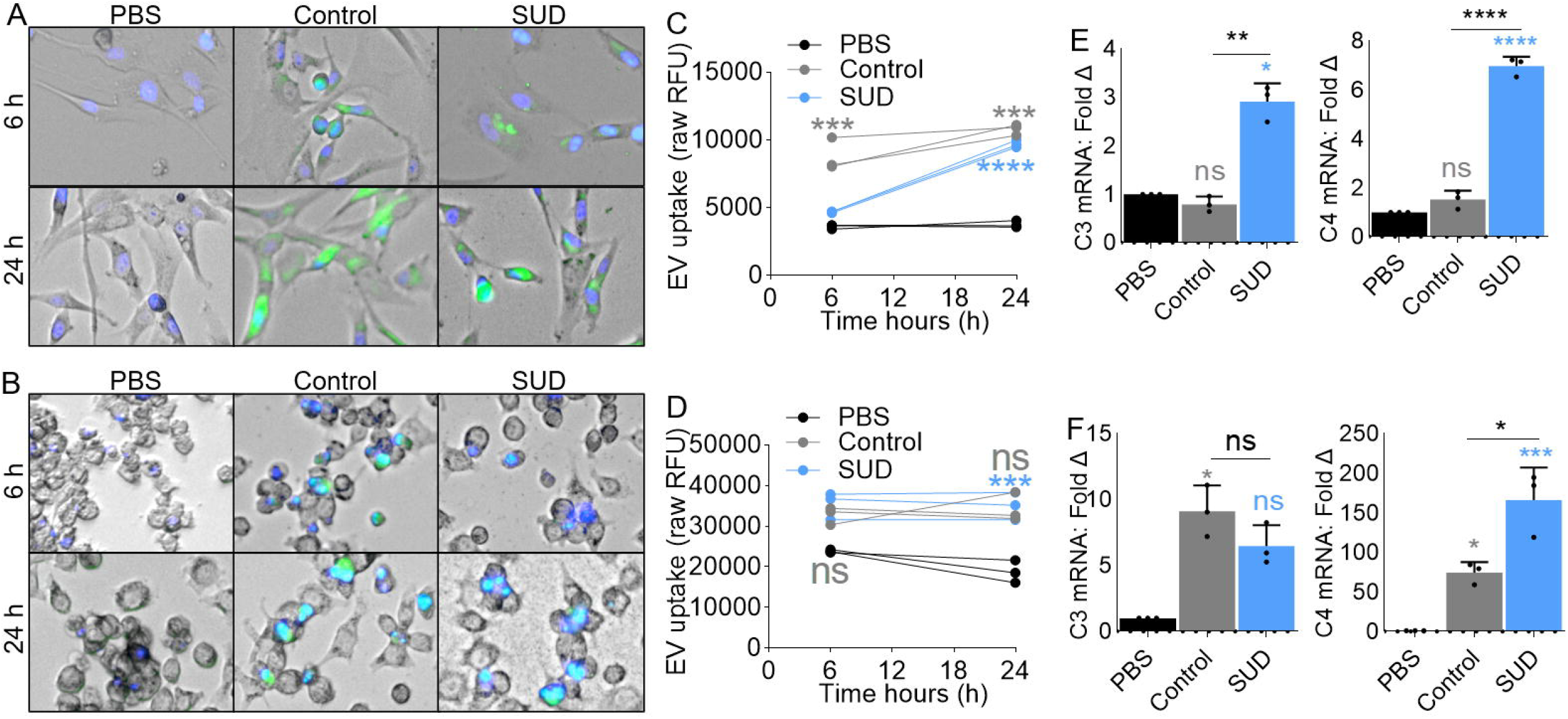
BA9 EVs are taken up by glia cells and the EVs altered glia cell gene expression. **A, B**) Representative 10x images of **A**) human primary astrocytes and **B**) HMC3 microglia incubated with SytoSELECT (green fluorescence) labeled BA9 EVs (50 μg) in the presence of NucBlue (a live cell stain, 30 μL/mL) and seeded in a glass-bottom 96 well plates. Images were taken at 6 hours and 24 hours using automated Lionheart FX microscope (Biotek). **C, D**) Uptake of BA9 EVs by **C**) astrocytes and **D**) microglia was analyzed by quantifying 3 fields of view with Gen5 software and presented as raw relative florescence units (RFU). **E, F**) Gene expression of **E**) human primary astrocytes and **F**) HMC3 microglia cells treated with control and SUD BA9 EVs was assessed by RT-qPCR. Cells treated with PBS was used as negative control. For panels C and D, statistical significance was determined by two-way ANOVA. **** p < 0.0001, *** p 0.0002 - 0.0009, ns = non-significant. For panels E and F, statistical significance was determined by ordinary one-way ANOVA (Šídák’s multiple comparisons test) and unpaired t test with Welch’s correction. **** p < 0.0001, *** p 0.0002 - 0.0009, ** p 0.0042, * p 0.0247, ns = non-significant.

### EVs reprogram the transcriptome of glia states

Since the protein content of SUD EVs, especially complement proteins C3 and C4A, were markedly different from that of control EVs, we sought to assess the effect of the EVs on glia cell transcriptome with a focus on C3 and C4A. In astrocytes, EVs from SUD but not controls significantly increased the levels of C3 and C4A mRNA after 24 hour treatment (**Figure 5E**). In contrast, both control and SUD EVs significantly increased microglia C3 and C4A mRNA (**Figure 5F**).

### EVs from SUD individuals promote collective migration of microglia cells

Since PIP2 is enriched in SUD EVs (**Figure 2F**) and PIP2, along with its phosphorylated product PIP3, plays crucial roles in maintaining cell polarity and polarized trafficking of molecules in cells, we investigated the role of EVs from control and SUD individuals in directional cell migration. Using our previously described wound healing assay^52^, we examined the migration of human microglia cells in response to a mechanical scratch wound, either in the absence (PBS) or presence of EVs isolated from control or SUD individuals. Representative images of the scratch areas at 0, 18, and 24 hours are shown in **Figure 6A**. Cells treated with PBS had approximately 89.9% and 89.5% wound area remaining at 18 and 24 hours, respectively (**Figure 6A, 6B, left**). In contrast, cells treated with control EVs exhibited 93.7% and 84.3% wound area at the same time points (**Figure 6A, 6B, middle**), indicating that EVs promote microglial migration. Notably, in the presence of EVs from SUD individuals, microglia cells showed even more enhanced migration, with 80.8% and 64.3% wound area remaining at 18 and 24 hours, respectively (**Figure 6A, 6B, right**). These findings suggest that EVs from SUD individuals significantly potentiate microglial migration beyond the levels induced by control EVs from the dlPFC.

**Figure 6:**
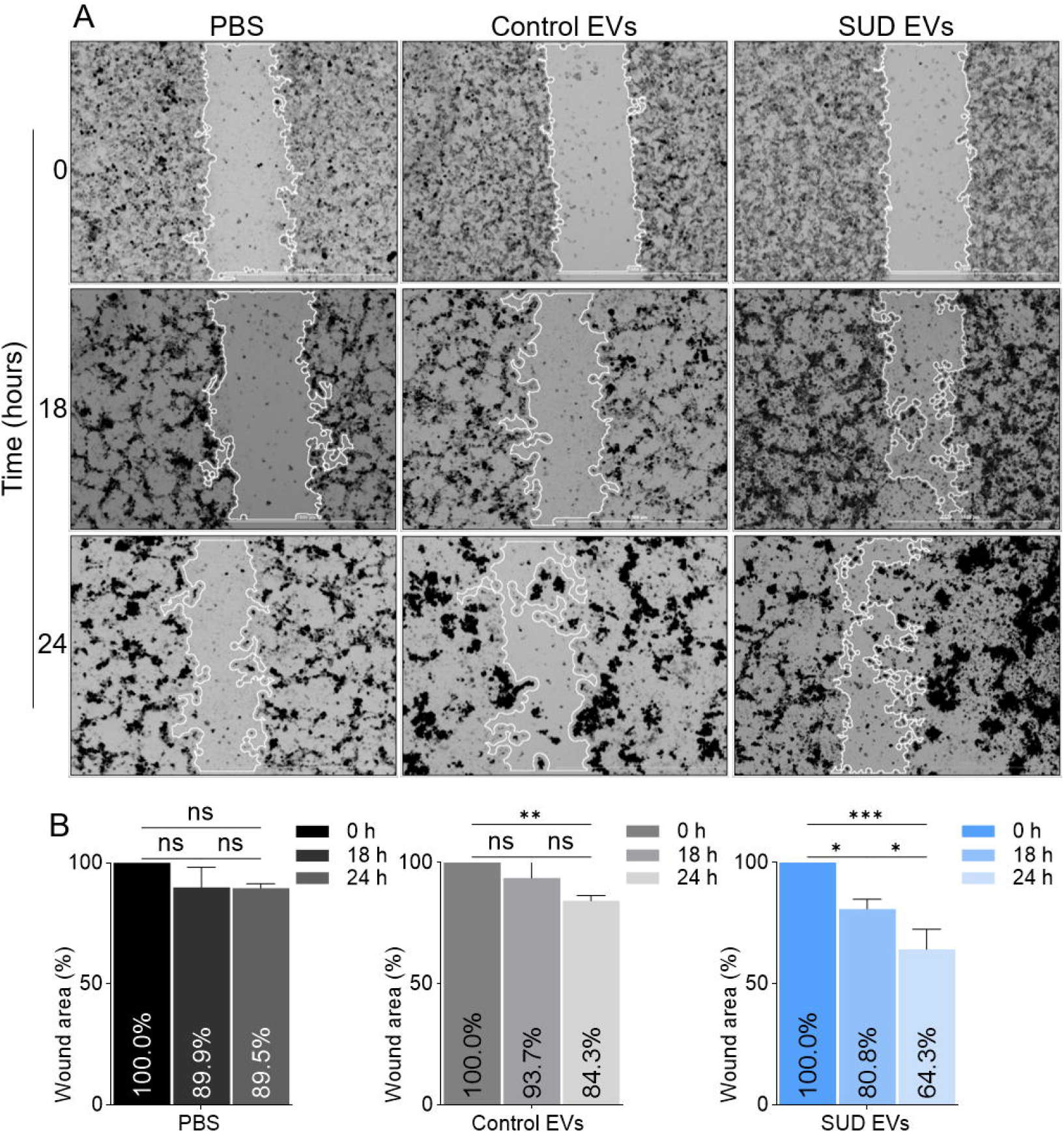
BA9 EVs regulate microglia cell migration. **A**) Representative images of wound tracks and areas. The vertical white lines track the wound areas. Images were capture with LionHeart with the 4X objective. Scale = 3000 μm. (A) Time and concentration dependent plot of wound area analyzed by quantifying 3 fields of view with Gen5 software and presented as percent wound area. Statistical differences were assessed by Ordinary oneway ANOVA with Tukey’s multiple comparisons test for panel C, and ordinary one-way ANOVA with Šídák’s multiple comparisons test and unpaired t test with Welch’s correction. *** p 0.0005, ** p 0.0088, * p 0.018 – 0.03, ns = non-significant.

## Discussion

In this study, we successfully isolated EVs from human postmortem brain tissue and elucidated changes occurring within the dlPFC in individuals with SUD. To our knowledge, our paired lipidomics and proteomics analyses of BA9 EVs are unprecedented in the field of SUD research. Previous studies, including ours, have primarily focused on the protein and nucleic acid composition of EVs, providing valuable insights on the roles that EVs play in human health and disease ^15, 21, 22^. However, our current comprehensive analysis represents a significant advancement in understanding the impact of SUD on the brain and the role of EVs in this process.

We detected relatively high numbers of total sphingolipid and phospholipid species in BA9 EVs, with an equal distribution between control and SUD groups. These findings align with existing literature showing that EVs are enriched in sphingolipids (sphingomyelin and ceramide) and glycerophospholipids containing saturated fatty acids ^56-58^. Prior studies have suggested that the detection of large numbers of sphingolipids in EVs may be due to contamination by lipoproteins or lipid droplets ^59^. However, our use of PPLC effectively separates EVs from other extracellular particles, countering this assumption ^26, 32^. Our analysis revealed that phosphoinositides, particularly phosphatidylinositol bisphosphate (PIP2), were abundant in BA9 EVs. Although total phosphoinositides did not significantly differ between control and SUD EVs, PIP2 was notably elevated in SUD EVs. PIP2 plays a critical role in maintaining cell polarity, directional cell migration, immune signaling, and inflammation ^60^. The significant elevation of PIP2 in SUD EVs suggests functional implications that merit further investigation.

Membrane lipids interact with proteins in various ways, potentially stabilizing the membrane environment for proteins to embed within EV membranes. We observed SUD-specific protein enrichment in BA9 EVs. The presence of tetraspanins (CD9, CD63, CD81) correlates with previous studies ^7, 32, 55, 61^. Our protein enrichment analysis indicated that SUD EVs may be pathologic, potentially activating various diseases, including infectious, neurological, and psychological disorders, as well as organismal injury and abnormalities. Of particular interest, SUD EVs exhibited elevated levels of C3 and C4A/4B complement proteins. The complement system is part of the innate immune system’s first line of defense against pathogens and plays a crucial role in neuronal health ^62^. However, overactivation of the complement cascade in the brain can lead to various neurological dysfunctions ^63, 64^, including autism spectrum disorder (ASD) ^65, 66^. While C4A has been implicated in regulating blood-brain barrier permeability ^67^ and the pathophysiology of psychiatric disorders such as schizophrenia ^68, 69^, our findings are novel in implicating C3 and C4A complement proteins in SUD.

Our flow cytometric analysis identified a higher number of microglia-derived EVs in SUD BA9 compared to controls. This finding suggests that SUD dysregulates the release of EVs in a cell type-dependent manner and that EVs derived from specific cellular origins in the brain, such as microglia or astrocytes, may have biological effects on bystander cells. The transfer of EV cargos, including biologically active nucleic acids, lipids, lipid metabolites, and proteins, has emerged as a mechanism by which cells alter biological processes within target cells and tissues, including the brain ^21, 33, 34, 61^. This aligns with our findings that BA9 EVs altered the levels of C3 and C4A/4B mRNA in astrocytes and microglia. Additionally, BA9 EVs from control individuals increased microglial migration, while EVs from SUD individuals further enhanced this migration. The increased migration in the presence of BA9 EVs indicates that these EVs are functionally bioactive, and SUD enhances their activity. Although the precise mechanisms by which SUD BA9 EVs potentiate cell migration remain unclear, the significant increase in wound closure (directional cell migration toward the wound area) in cells treated with SUD EVs supports our lipidomic data, which showed PIP2 enrichment in SUD EVs. Given the increasing evidence that migration is crucial in cell-cell communication and interactions with the microenvironment as reviewed by Miskolci *et al*., 2021 ^70^, and therefore important in physiological and pathological processes such as wound healing, immune response, pathogen dissemination, and neuroinflammation, our findings warrant further investigation into the role of BA9-derived EVs in SUD pathophysiology.

Neuroinflammation is a key component in the pathogenesis of numerous CNS diseases and is thought to contribute to neural adaptations following chronic drug exposure ^71^. Neuroinflammation includes gliosis, microglial activation, and the release of cytokines, chemokines, and pro-inflammatory factors, which are all implicated in the brain’s response to chronic substance use ^72-74^. Furthermore, increased presence of microglial-derived EVs in SUD (**Figure 7**) support the notion that drug use heightens neuroinflammatory processes, potentially leading to neuropathology. Our findings that BA9 EVs alter the levels of C3 and C4A/4B mRNA in astrocytes suggest that EVs could play a role in astrocyte-mediated neuroinflammation and synaptic modulation. This is consistent with studies showing that astrocytes play a critical role in the uptake of synaptically-released glutamate and are affected by the activity levels of dopamine (DA) neurons, which are often dysregulated in SUD ^75^. Drug-induced dysregulation of neuroimmune signaling may compromise neuronal function, exacerbate neurodegeneration, and increase neurotoxicity, all of which can contribute to drug-related behaviors through microglia and other glia-mediated synaptic remodeling ^74^. The dysregulation of EV cargo in SUD, particularly the enrichment of PIP2 (**Figure 2**) and complement proteins C3 and C4A/4B (**Figure 5**), points to specific pathways that may be targeted to alleviate the neuroinflammatory and neurodegenerative consequences of SUD. PIP2’s role in cell signaling, immune responses, and maintaining cell polarity ^76^ underlines its potential impact on neuronal health and the progression of neuroinflammation in SUD. Similarly, the involvement of complement proteins in SUD highlights a possible mechanistic link to the broader neuroimmune response observed in these individuals. The overactivation of these pathways in SUD could lead to detrimental effects on synaptic integrity and cognitive function.

Overall, our study underscores the complex interplay between EVs, glial cells, and neuroinflammation in SUD. The identification of specific lipidomic and proteomic changes in EVs from SUD individuals provides valuable insights into the molecular mechanisms underlying the disease. These findings highlight potential biomarkers and therapeutic targets for addressing the neuroinflammatory and neurodegenerative effects of chronic substance use. Future research should focus on elucidating the precise roles of these EV components in mediating neuroimmune responses and their potential as targets for intervention in SUD.

## Limitations

Although this study presents groundbreaking observations of the possible functions of BA9 EVs using human postmortem brain tissues, there are limitations. Because of the small sample size, the investigation of specific SUDs – CUD and OUD - was not powered and these could not be analyzed independent of each other. Likewise, sex and ancestry differences were not analyzed due to the limited sample size. Future studies are needed to investigate the contributions of specific substances of abuse, sex, and race to the BA9 EVs composition identified in this study.

## Conclusions

Our study demonstrates the effective use of novel PPLC technology to isolate BA9 extracellular vesicles (EVs) and elucidate their lipidomic and proteomic profiles in the brains of individuals with SUD as compared to individuals without SUD. We identified significant alterations in EV secretion, uptake by glial cells, and their impact on glial transcriptome and collective migration. These findings contribute to a deeper understanding of the role of brain-derived EVs in the context of SUD, highlighting their potential involvement in pathogenic processes and their utility as biomarkers for SUD diagnosis. As the field progresses towards a more comprehensive and integrated analysis of brain-derived EVs, our study underscores the importance of examining these vesicles in understanding the molecular mechanisms of SUD. Future research should aim to determine whether the observed alterations in BA9-derived EVs are also present in peripheral EVs, such as those derived from blood, or in other brain regions. This will be crucial for validating EVs as reliable biomarkers and therapeutic targets for SUD.

## Declarations

## Ethical Approval and Consent to participate

Not applicable.

## Consent for publication

All authors read and approved the publication of this manuscript.

## Competing interests

The authors report no biomedical financial interests or potential conflicts of interest.

## Availability of data and materials

The lipidomics and proteomics datasets are included within the article and its additional files.

## Funding

This work was supported by National Institutes of Health funding (Grant No. R01DA042348 [to CMO]; Grant Nos. R01DA050169 and R21/R33DA053643 [to CMO]; and Grant Nos. R01DA044859 [to CWB and RGO]; Independent Research Fund Denmark *3166-00139B* [to RGO]; The John S. Dunn Foundation to CWB

## Author contributions

CMO, RGO, and CWB conceptualized the study. CWB provided brain tissue. RGO WN, CG, YS, CV AN, AK conducted the experiments. WN, BCO, CG, YS, CV, AN, AK, RGO, CWB, and CMO conducted data analyses. WN, BCO, CG, YS, CV, AN, AK, RGO, CWB, and CMO wrote the original draft of the paper, provided text, reviewed and edited the manuscript. CMO, RGO, and CWB wrote the final manuscript.

## Acknowledgments

This work was supported, in part, by the New York Medical College Histology and Imaging Core, Iowa Hybridoma Bank to CMO. CWB and RGO were supported by NIDA R01DA044859. RGO was supported by Independent Research Fund Denmark Agency grant 3166-00139B. Drs. Vary, Gartner and Santos-Ortega and the MHIR Proteomics and Lipidomics Core Facility are supported by NIH/NIGMS award 1P20GM12130 to Lucy Liaw. Dr. Vary and his Core facility also supported under NIH/NIGMS award U54GM115516 (C. Rosen and G. Stein, principal investigators). We are grateful for the invaluable donations and participation from families, as well as for the generous collaboration of the medical examiners at the Harris County Institute of Forensic Sciences. TEM images were generated by Tamara Howard at the University of New Mexico HSC-Electron Microscopy Facility, which is supported by The University of New Mexico Health Sciences Center.

## Figure legends

**Supplemental Figure 1:** Lipid Ontology (LION) enrichment analysis presented as heatmap for **A**) 98 positive and **B**) 385 negative polarity lipid species. **C**) LION-PCA and LION-terms for positive (Top) and negative (Bottom) polar lipids.

**Supplemental Figure 2: A**) PC plot of BA9 EVs. **B**) Clusters of dysregulated proteins identified in BA9 EVs. **C**) Canonical pathway analysis. **D**) Merged IPA networks (5 networks).

## Notes

### Competing Interest Statement

The authors have declared no competing interest.

