## Supplementary figures and images for "Lipidomic and Proteomic Insights from Extracellular Vesicles in Postmortem Dorsolateral Prefrontal Cortex Reveal Substance Use Disorder-Induced Brain Changes"

### Supplemental Figure 1

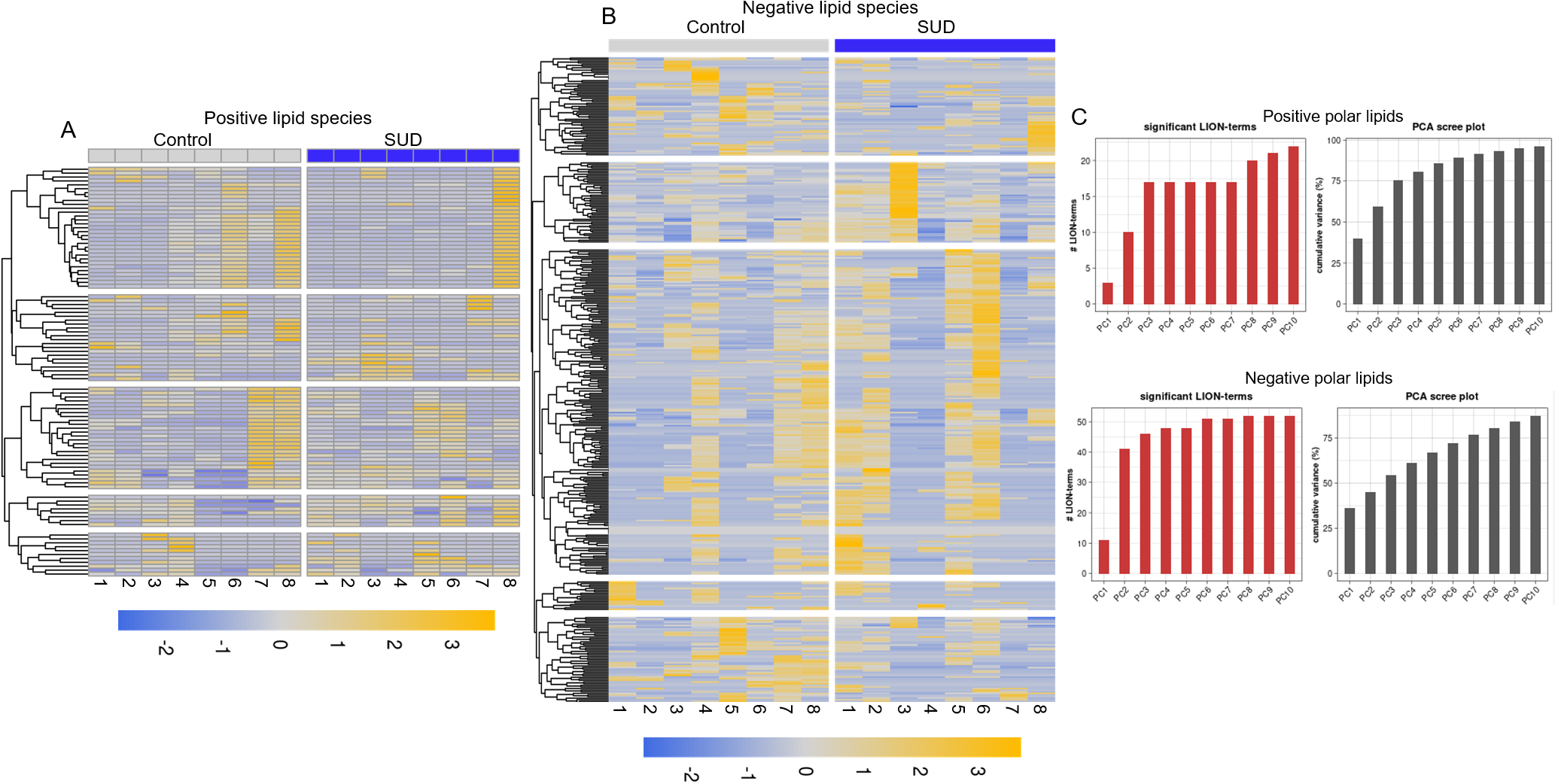

### Supplemental Figure 2

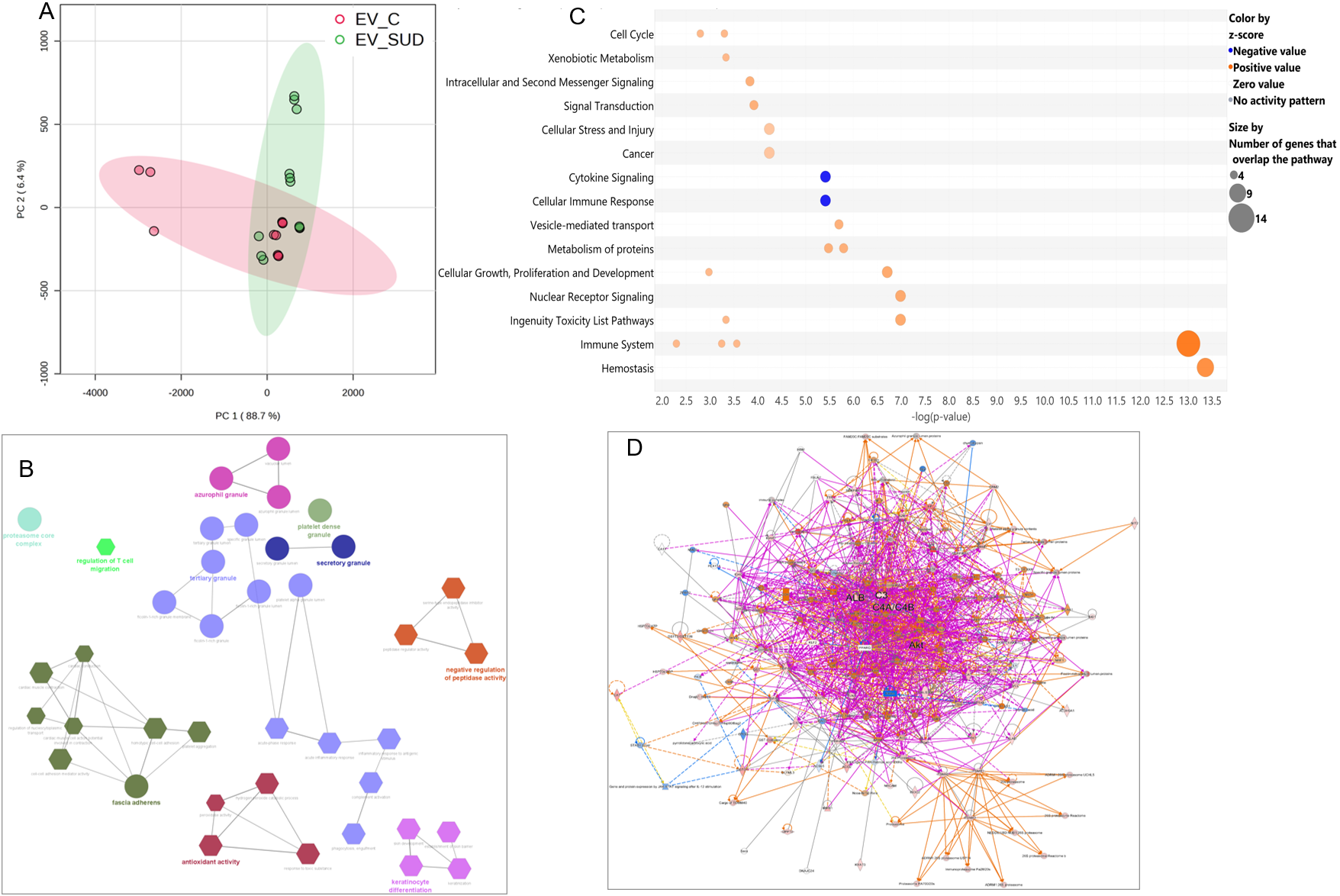
